# DEER-PREdict: Software for Efficient Calculation of Spin-Labeling EPR and NMR Data from Conformational Ensembles

**DOI:** 10.1101/2020.08.09.243030

**Authors:** Giulio Tesei, João M. Martins, Micha B. A. Kunze, Yong Wang, Ramon Crehuet, Kresten Lindorff-Larsen

## Abstract

Owing to their plasticity, intrinsically disordered and multidomain proteins require descriptions based on multiple conformations, thus calling for techniques and analysis tools that are capable of dealing with conformational ensembles rather than a single protein structure. Here, we introduce DEER-PREdict, a software to predict Double Electron-Electron Resonance distance distributions as well as Paramagnetic Relaxation Enhancement rates from ensembles of protein conformations. DEER-PREdict uses an established rotamer library approach to describe the paramagnetic probes which are bound covalently to the protein. DEER-PREdict has been designed to operate efficiently on large conformational ensembles, such as those generated by molecular dynamics simulation, to facilitate the validation or refinement of molecular models as well as the interpretation of experimental data. The performance and accuracy of the software is demonstrated with experimentally characterized protein systems: HIV-1 protease, T4 Lysozyme and Acyl-CoA-binding protein. DEER-PREdict is open source (GPLv3) and available at github.com/KULL-Centre/DEERpredict and as a Python PyPI package pypi.org/project/DEERPREdict.

## Introduction

A detailed understanding of protein function often requires an accurate description of the structure and dynamics of a protein. The characterization of protein complexes as well as multi-domain and disordered proteins is typically achieved by combining experimental techniques of distinct spatial resolution [1]. Among the many different experimental techniques that may be used, we focus here on (i) a pulsed electron paramagnetic resonance (EPR) technique called double electron-electron resonance (DEER) and (ii) a nuclear magnetic resonance (NMR) method called paramagnetic relaxation enhancement (PRE). While the two methods differ substantially in their physics and applications, they have in common that they generally involve adding so-called spin-labels to the protein of interest.

DEER, also sometimes known as pulsed electron-electron double resonance (PELDOR), [2–6] relies on probing magnetic dipole-dipole interactions that are sensitive to distributions of residue-residue distances ranging from ∼ 1.8 nm to ∼ 8 nm, and up to 16 nm in deuterated soluble proteins [7–10]. For proteins, DEER generally requires site-directed spin labeling (SDSL) to functionalize a pair of selected residues with paramagnetic probes, e.g. 1-Oxyl-2,2,5,5-tetramethylpyrroline-3-methyl methanethiosulfonate (MTSSL) [4].

PRE NMR also makes use of SDSL to provide information on the average proximity of protein backbone nuclei up to ∼ 3.5 nm away from the unpaired electron of the paramagnetic probe [11]. The dependence of the rate of relaxation enhancement on the electron-proton distance, *r*, scales as ⟨*r*^−6^⟩, making the measurement particularly sensitive to contributions from different probe conformations [11].

Since spin labels are conformationally dynamic, both protein and paramagnetic probes need to be described by conformational ensembles to obtain accurate predictions of DEER and PRE observables from molecular models [12–14]. Molecular dynamics (MD) simulations are one approach to obtain conformational ensembles that model the structure and dynamics of spin-labels for the calculation of EPR and NMR data [15–18]. While such analyses can provide unique insight into the motions of and interactions between protein and spin-label [19], they may be relatively expensive computationally. Further, many studies integrate results from multiple probe positions, or pairs thereof, which may be difficult to represent in a single MD simulation with explicit representations of the probes.

Another approach is to use conformational analysis of the spin-label combined with modelling of the dynamics [20–23]. Such analyses suggest that the conformational variation of spin-labelled sites is rotameric, i.e. it can be relatively well described by a finite number of defined structures. Thus, in the calculation of DEER data, rapid modeling of dynamic paramagnetic probes was made possible with the introduction of the rotamer library approach (RLA) applied to the MTSSL probe by Polyhach *et al*. [24].

Here, building and expanding on earlier work [3, 24–27], we developed a software tool for fast predictions of DEER and PRE observables from large conformational ensembles using the RLA. We present our implementation, distributed as the DEER-PREdict software, and test it against experimental data on HIV-1 Protease, T4 Lysozyme and the Acyl-CoA-Binding Protein. This software has been previously used for the calculation of both intra- and intermolecular DEER and PRE NMR data [28, 29], and has some overlap with the features in RotamerConvolveMD [25] (github.com/MDAnalysis/RotamerConvolveMD). DEER-PREdict is open-source, documented (deerpredict.readthedocs.io) and open to contributions from the community.

## Design and Implementation

DEER-PREdict is written in Python and is available as a Python API, which facilitates its integration within larger data pipelines. Predictions of DEER and PRE data are carried out via the DEERpredict and PREpredict classes. Both classes are initialized with protein structures (provided as MDAnalysis [30] Universe objects) and spin-labeled positions (residue numbers and chain IDs). As shown in the *Results* section, the calculations are triggered by the *run* function, which also sets additional attributes such as the paths of input and output files as well as experiment-specific parameters. Per-frame data is saved in compressed binary files (HDF5 and pickle files) to allow for fast calculations of ensemble averages in reweighting schemes.

For the presented software, we adopt a procedure of rotamer placement and evaluation of labeled sites which is analogous to the RLA of Polyhach *et al*. [24], and we build on this previous work to implement fast calculations of DEER and PRE observables from large structural ensembles, such as MD trajectories.

### Rotamer Library Approach

Rotamer libraries have a long history in protein structural analysis [31], with an early application being to study side-chain packing [32]. Several other applications of this approach were later employed, e.g. in homology modeling and protein design [33, 34]. In our implementation, the RLA is used to insert the rotamer conformations of a paramagnetic probe at the spin-labeled site and to calculate the Boltzmann weight of each conformer. By default, we use the MTSSL 175 K rotamer library by Polyhach et al. [24], which was filtered to include only the *χ*_1_*χ*_2_ conformations that are most commonly found in crystal structures of T4 Lysozyme [35]. As shown by Klose *et al*. [26], this selection criterion increases the accuracy of the calculated electron-electron distance distributions. The code is, however, general and it is possible to add new rotamer libraries by providing a text file containing the Boltzmann weights of each rotamer state 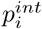, a topology file (PDB format) and a trajectory file (DCD format) where rotamers are aligned with respect to the the plane defined by C*α* atom and C–N peptide bond. These files should be included in the *lib* folder and listed in the yaml file *DEERPREdict/lib/libraries*.*yml*. The default MTSSL 298 K MC/UFF C*α*S*δ* rotamer libraries of the Matlab-based MMM modeling toolbox [13] are also provided in the DEER-PREdict package.

Following the alignment of the rotamer to the protein backbone (C*α*, C and N atoms), the calculation of the Boltzmann weights is based on the sum of internal, 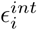, and external, 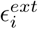, energy contributions. The internal contribution is taken from Polyhach et al. [24] and results from the clustering of representative dihedral combinations from MD simulations. The normalized frequency of each cluster throughout the MD trajectory was used to determine the Boltzmann probability, 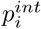, of a given *i*^*th*^ state, which readily can be converted into an internal energy contribution, 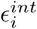, via Boltzmann inversion. On the other hand, the external energy contribution is calculated on the fly as the dispersion interaction energy between heavy atoms of rotamer and protein residues within a 1-nm cutoff, using the pairwise 6-12 Lennard-Jones potential of the CHARMM36 force field.

The overall probability of the *i*^*th*^ rotamer state is then calculated as

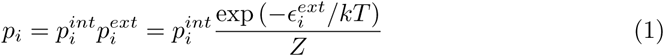

where 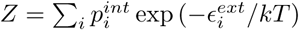 is the steric partition function quantifying the fit of the rotamer in the embedding protein conformation. Low values of *Z* result from large probe-protein van der Waals interaction energies, suggesting a tight placement of the spin label either due to a displacement of the rotamers or indicative of a wild-type conformation made inaccessible by the presence of the MTSSL probe. Especially in folded proteins, probes located in closely packed regions are likely to induce changes in the ensemble of the spin-labeled protein compared to the native form, and should be avoided in designing SDSL experiments. Therefore, in the calculation of DEER or PRE NMR observables, frames with *Z* < 0.05 are discarded to preclude spurious conformers from contributing to the ensemble average [24]. For the MTSSL 175 K rotamer library, a *Z* cutoff of 0.05 is compatible with distributions of 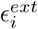 values where at most one of the 46 rotamers has 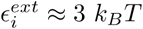 while the rest has 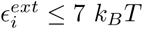. We observed that the results shown in this paper are virtually insensitive to the choice of the *Z* cutoff between 0.05 and 0.5 (see Figure S1), therefore, in DEER-PREdict the default *Z* cutoff can be conveniently replaced by a user-provided value.

### Predicting the DEER signal from structural ensembles

Electron-electron distance distributions extracted from DEER experiments, e.g. using the DeerLab package [36], have previously routinely been compared with distributions predicted using the RLA implemented in the Matlab-based MMM modeling toolbox (http://www.epr.ethz.ch/software) [13]. Since MMM intrinsically operates on single structures, we and others had to resort to wrapper scripts to compute distance distributions of large ensembles, such as MD trajectories [3, 25, 37]. With the program presented herein, we provide a tool to conveniently predict DEER distance distributions from large conformational ensembles, which can be easily integrated in reweighting schemes such as the Bayesian/maximum entropy procedure [1, 14, 38, 39].

For each trajectory frame or conformation of a given ensemble, the rotamers from the library are placed at the spin-labeled position (Figure 1A) and the distances between all pair combinations of N-O paramagnetic centers are calculated. The resulting matrix of pair-wise distances is then used to compute the distance distribution weighted by the combined probability of each probe conformation, *p*_*i*_ × *p*_*j*_, with *p*_*i*_ and *p*_*j*_ being the conformation probabilities of rotamers *i* and *j*. After averaging over all the frames, a low-pass filter is applied to the distance distribution for noise reduction [40],

**Fig 1.**
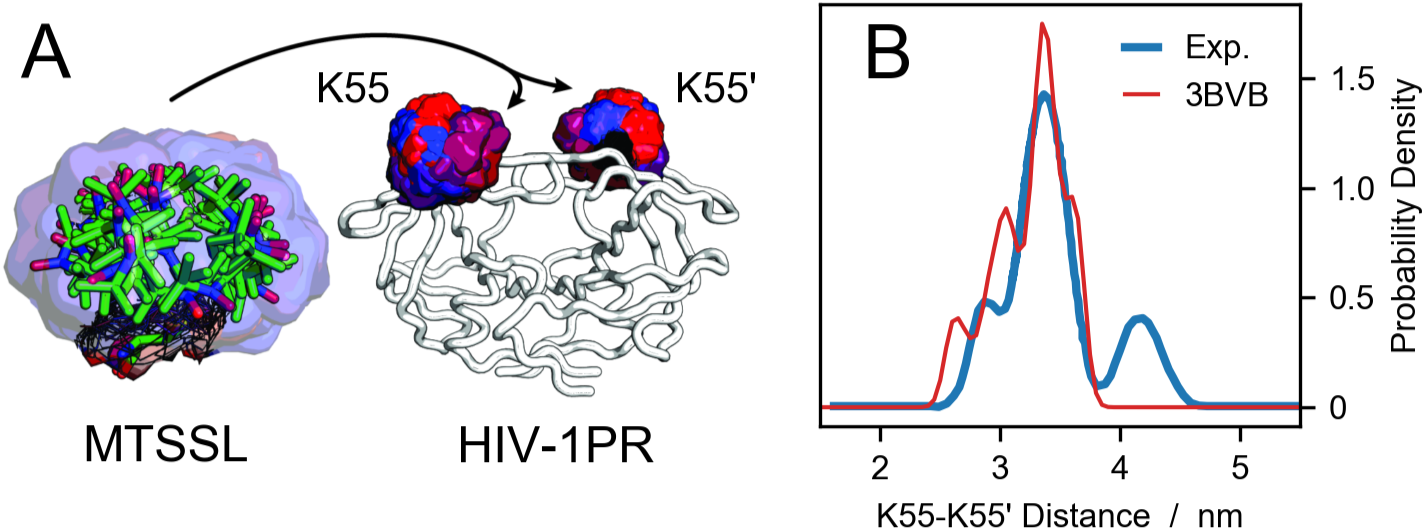
Probe placement scheme and comparison to DEER data. (A) A pool of 46 conformations of the MTSSL probe from the rotamer library are aligned to the backbone of residues K55 and K55’ of HIV-1 protease. The color code represent the Boltzmann weights of each rotamer, increasing from blue to red. (B) Electron-electron distance distribution for HIV-1 protease spin labeled at residues K55 and K55’. The blue line is the experimental data from Torbeev *et al*. [44] whereas the red line is the prediction using DEER-PREdict and a crystal structure of HIV-1 protease (PDB code 3BVB).

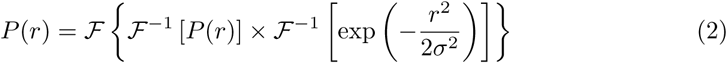

where ℱ and ℱ^−1^ are the Fourier transform and inverse Fourier transform operators, respectively, whereas *σ* is the standard deviation of the low-pass filter. The resulting *P* (*r*) is a smooth curve even for the analysis of a single protein conformation (Figure 1B). The standard deviation of the low-pass filter can readily be provided by the user through the option *filter_stdev* of the *run* function in the *DEERpredict* class, overriding the default value of 0.5 Å. The average over the trajectory frames can be weighted by a user-specified list of weights e.g. to remove the bias from enhanced sampling simulations.

The dipolar modulation signal can be back-calculated from the distance distribution, *P* (*r*), via the following integral [41]

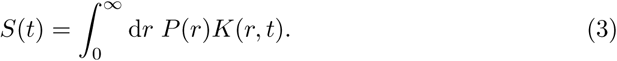

*K*(*r, t*) is the DEER kernel

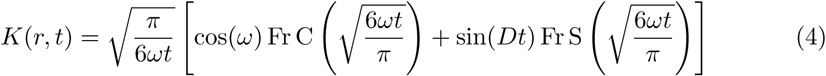

where FrC and FrS are Fresnel cosine and sine functions, and *ω* is the dipolar frequency

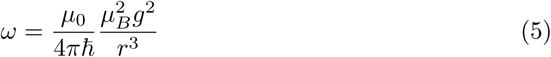

where *μ*_0_ is the permeability of free space, *μ*_*B*_ is the Bohr magneton and *g* is the electron g-factor. The inter-probe distances and the time range from [0, *r*_*max*_] and [*t*_*min*_, *t*_*max*_] with increments d*r* = 0.05 nm and d*t*, respectively. The default values *r*_*max*_ = 12 nm, *t*_*min*_ = 0.01 *μ*s, *t*_*max*_ = 5.5 *μ*s and d*t* = 0.01 *μ*s can be overridden by the user.

Following the correction of the experimental DEER time trace for the intermolecular background [36, 42], the resulting form factor can directly be compared with

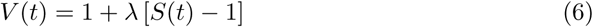

where 0.02 ≤ *λ* ≤ 0.5 is the modulation depth of the experimental signal [43], quantifying the efficiency of the DEER pump pulse [8].

### Prediction of PRE rates and intensity ratios

In analogy to the calculations of electron-electron distances to predict DEER distributions, we extended the use of the RLA to electron-proton separations to improve the accuracy of PRE predictions. We focus here is on PRE NMR experiments that probe the increase in transverse relaxation rates of any backbone proton due to the dipolar interaction with the unpaired electron of the paramagnetic probe:

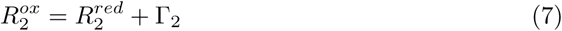

where 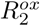 and 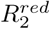 are the transverse relaxation rates in the presence of the spin label in the oxidized or reduced (diamagnetic) state, respectively. We note that it is also possible to measure PREs on other atoms and to probe longitudinal relaxation enhancement, and it would be possible to include such measurements in future versions of DEER-PREdict.

A description of the enhancement of the transverse relaxation due to dipole-dipole interactions in paramagnetic solutions was first proposed by Solomon and Bloembergen [45, 46]

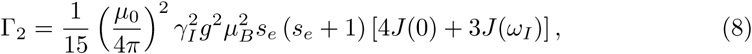

where *γ*_*I*_ and *ω*_*I*_ are the gyromagnetic ratio and the Larmor frequency of the proton, respectively, whereas *s*_*e*_ is the electron spin quantum number, equal to 1/2 for nitroxide probe systems. The spectral density function *J* (*ω*_*I*_) can be described using a model-free formalism [47–50], which takes into account the overall molecular tumbling in the external magnetic field as well as the internal motion of the spin label:

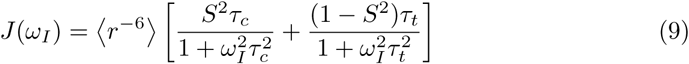

where

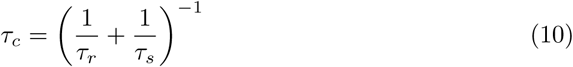

and

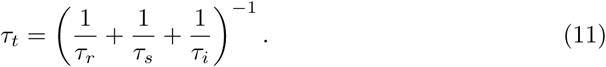

*τ*_*r*_ is the rotational correlation time of the protein, *τ*_*s*_ is the effective electron correlation rate and *τ*_*i*_ is the correlation time of the internal motion (effective correlation time of the spin label). For MTSSL probes, *τ*_*s*_ ≫ *τ*_*r*_ and *τ*_*c*_ ≈ *τ*_*r*_ [51]. The value of *τ*_*c*_ depends on protein size and structure and is generally of the order of 1–10 ns [27, 52–55]. For *τ*_*i*_, values between 100 to 500 ps can be assumed, based on e.g. ^15^N spin relaxation rates and MD simulations [56, 57]. In general, *τ*_*c*_ and *τ*_*i*_ can be specified as user input in DEER-PREdict.

For the generalized order parameter, *S*, we use the factorization into contributions from radial and angular internal motions introduced by Brüschweiler *et al*. [49], 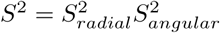. The expressions for 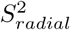 and 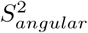 were derived from a jump model that treats the *N* conformers of the rotamer library as *N* discrete states with equal probabilities (1*/N*) [50]. In reality, the various dihedral angles of the spin label have different free energy barriers, resulting in residence times between jumps ranging from less than 1 to several ns [17].

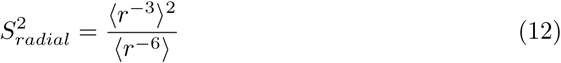

where *r* is the proton-electron distance and the brackets denote averages over the conformers weighted by the respective Boltzmann weights, *p*_*i*_, i.e. 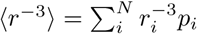 and 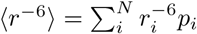.

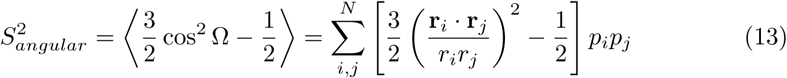

where Ω is the angle between the vectors **r**_**i**_ and **r**_**j**_, connecting a backbone proton with the *i*th and *j*th rotamer states, respectively. The relaxation enhancement rate for a single protein structure is calculated using Equation 8, and assuming that the motion of the paramagnetic label is much faster than the protein conformational changes, the ensemble average is estimated as

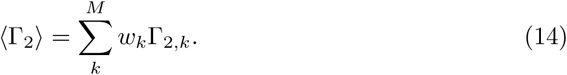

where *M* is the number of configurations or frames of the simulation trajectory. In the case of unbiased simulations, the statistical weights, *w*_*l*_, are simply 1*/M*. Optionally, a list of weights can be provided by the user, e.g. to reweight a biased MD simulation [58, 59] or to incorporate the prediction of the PRE rates into a Bayesian/maximum entropy reweighting scheme [1].

For samples with particularly high PRE rates it can be infeasible to obtain G_2_ from multiple time-point measurements [60]. In such and other cases, the PRE is sometimes probed indirectly from the ratio of the peak intensities in ^1^H,^15^N-HSQC spectra of the spin-labeled protein in the oxidized and reduced state. Assuming that the intensity of the proton magnetization decays exponentially — by transverse relaxation only — during the total INEPT time of the HSQC measurement [61], *t*_*d*_, the intensity ratio is estimated as

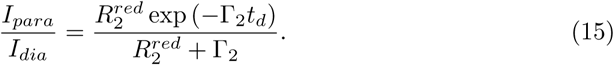

### Requirements and Installation

The main requirements are Python 3.6–3.8 and MDAnalysis 1.0 [30, 62]. In an environment with Python 3.6–3.8, DEER-PREdict can readily be installed through the package manager PIP by executing

1. `pip install DEERPREdict`

### Package Stability

Tests reproducing DEER and PRE data for the protein systems studied in this article, as well as for a nanodisc [29], are performed automatically using Travis CI (travis-ci.com/github/KULL-Centre/DEERpredict) every time the code is modified on the GitHub repository. The same tests can also be run locally using the test running tool pytest.

## Results

In the following, we present applications of our tool to the prediction of DEER distance distributions and PRE intensity ratios of three folded proteins.

The code snippets reported in this section pertain to DEER-PREdict version 0.1.4. A Jupyter Notebook to reproduce the results shown below (*article*.*ipynb*) can be found in the *tests/data* folder on the GitHub repository. The up-to-date documentation is available at deerpredict.readthedocs.io.

### Case study 1: DEER data for HIV-1 Protease

HIV-1 protease (HIV-1PR) is a homodimeric aspartic hydrolase involved in the cleavage of the gag-pol polyprotein complex. The inhibition of this process affects the life cycle of the HIV-1 virus, rendering it noninfectious [63]. The HIV-1PR monomer is composed of 99 residues and presents a structurally stable core region (residues 1-43 and 58-99) and a dynamic region characterized by a *β*-hairpin turn, called the flap (residues 44-57). The active site is located at the interstice between the core regions of the two monomers, in proximity to the catalytic D25 residues. This cavity is closed off by the dynamic flap regions, which are considered to act as a gate controlling the access to the active site. The dynamics of the flap regions are of utmost importance for the development of inhibitors, and have been extensively studied, both experimentally and *in silico* [44, 64–69]. Based on the relative position of the flaps, three main conformational states have been proposed. In X-ray crystallography, the closed state is typically observed for the ligand-bound enzyme (e.g. PDB codes 3BVB [70] and 2BPX [71]), the semi-open state is predominant for the apo form (e.g. PDB code 1HHP [72]) whereas the wide-open state has been observed for variants (e.g. PDB codes 1TW7 [73] and 1RPI [74]) [69]. In DEER measurements, these conformational states can be resolved by spin-labeling sites K55 and K55’ (see Figure S2).

To assess the predictive ability of DEER-PREdict, we generated conformational ensembles of the HIV-1PR homodimer via two different approaches: (a) a single 500-ns unbiased MD simulation, and (b) four independent 125-ns MD simulations restrained with experimental residual dipolar couplings (RDC) data [58, 75] from Roche et *al*. [65, 66] (see SI for methodological details). The initial configuration of our simulations is the X-ray crystal structure of the active-site mutant D25N bound to the inhibitor Darunavir (PDB code 3BVB).

Figure 2 presents a comparison of experimental DEER distance distributions and echo intensity curves with predictions from simulation trajectories of 1,000 frames sampled every 0.5 ns. The echo intensity curves are calculated using Equation 6, where the *λ* is estimated to 0.0922 by fitting the experimental dipolar evolution function to the corresponding curve derived from the experimental *P* (*r*) via Equation 3. For a single trajectory, the analysis is performed in 40 s on a 1.7 GHz processor by running the following code:

**Fig 2.**
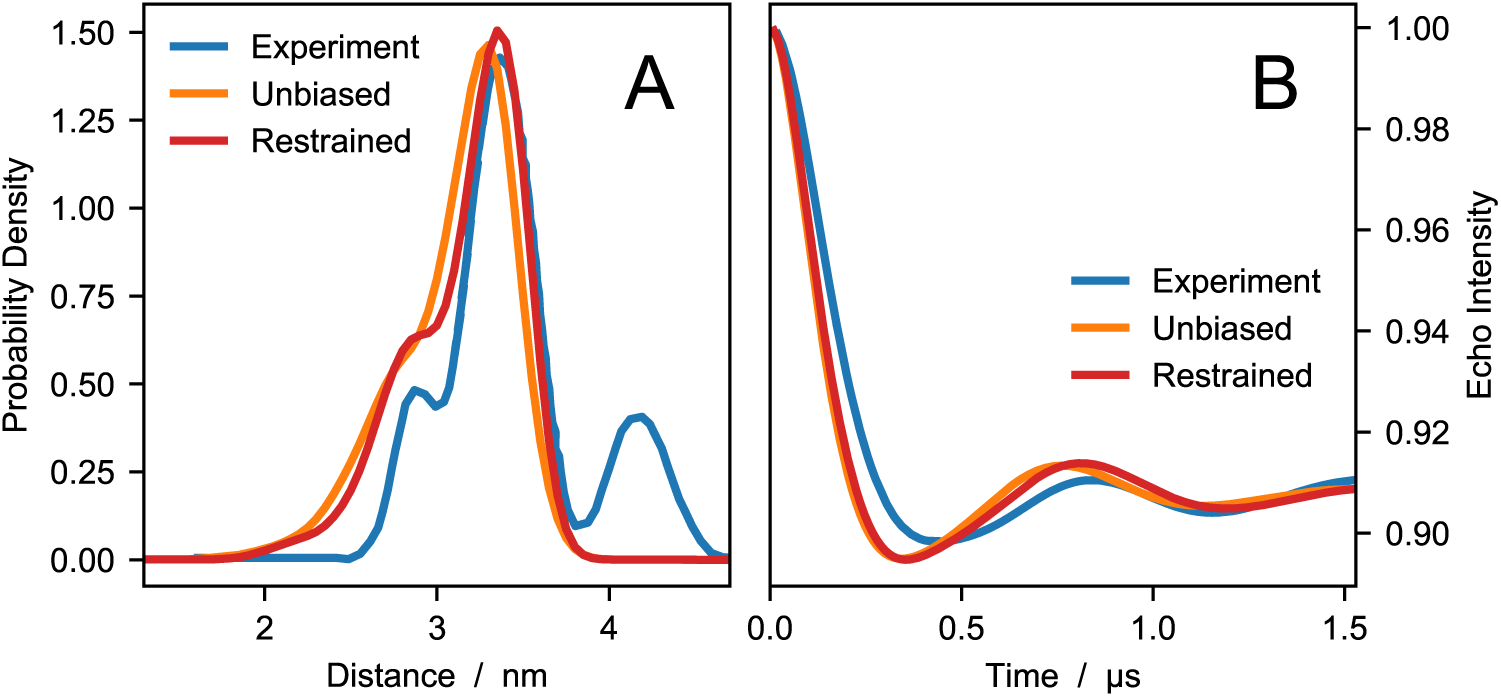
Comparing experiments and simulations for HIV1-PR. DEER distance distributions (A) and echo intensity curves (B) obtained by Torbeev *et al*. [44] from DEER experiments (blue), and calculated using DEER-PREdict from unbiased (orange) and RDC ensemble-biased MD simulations (red).

1. `import MDAnalysis`
2. `from DEERPREdict. DEER import DEERpredict`
3. `u = MDAnalysis. Universe (’ conf. pdb’,’ traj. xtc’)`
4. `DEER = DEERpredict (u, residues =[55, 55], chains =[’ A’,’B’], temperature = 298)`
5. `DEER. run ()`

The third line generates the MDAnalysis Universe object from an XTC trajectory and a PDB topology. The fourth line initializes the DEERpredict object with the spin-labeled residue numbers and the respective chain IDs. The fifth line runs the calculations and saves per-frame and ensemble-averaged data to *res-55-55*.*hdf5* and *res-55-55*.*dat*, respectively, as well as the steric partition functions of sites K55 and K55’ to the file *res-Z-55-55*.*dat*.

In the experimental distance distribution, the main peak at ∼ 3.3 nm corresponds to the closed state whereas the second peak between 4 and 5 nm is characteristic of the wide-open state. The shoulder peak at ∼ 2.8 nm has been identified as an open-like state known as the curled/tucked conformation [9, 76, 77]. The results of our unbiased and restrained simulations are in substantial agreement with the findings of Roche *et al*. [65, 66], indicating that the flaps of the inhibitor-free HIV-1PR are predominantly in closed conformation. Compared to the distance distribution calculated from the starting configuration of PDB code 3BVB (see Figure 1), predictions based on MD trajectories more accurately reproduce the shape of the shoulder and the main peak of the experimental *P* (*r*). Moreover, using the RDC data as restraints leads to a significant improvement in the agreement between simulations and experiments, with the RMSD decreasing from 0.07 for the unbiased to 0.03 for the RDC ensemble-biased simulations. However, in the simulations we do not observe the wide-open state. This discrepancy could be due to insufficient sampling or could be attributed to the difference in sequence between the simulated protein and the experimental construct.

### Case study 2: DEER data for T4 Lysozyme

Lysozyme from the T4 bacteriophage (T4L) has long been used as a model system in the study of protein structure and dynamics [78–83]. Here, we focus on the L99A and the triple L99A-G113A-R119P mutants which are structurally similar and mainly differ in the relative populations of their major conformational states. The L99A variant presents a 150 Å^3^ hydrophobic pocket capable of binding hydrophobic ligands and has been thoroughly studied to further our understanding of the dynamics and selectivity of the binding pocket [78, 84]. The L99A variant occupies two distinct conformational states: the ground state (G) and the transient excited state (E), amounting for 97% and 3% of the population, respectively. The large-scale motions converting the G into the E state occur on the millisecond time scale and result in the occlusion of the cavity, which is occupied by the side chain of F114 in the E state [82]. The additional G113A and R119P mutations in the triple-mutant variant interconvert the populations of the conformational states to 4% for the G state and 96% for the E state [82] — note that, here and in the following, we refer to the G and E states based on their structural similarity to the L99A variant rather than on their relative populations. These conformational equilibria have been studied by DEER for various pairs of spin-labeled sites, which effectively resolve the G and E states as separate peaks of the *P* (*r*) [83].

Here, we compare DEER distance distributions calculated with DEER-PREdict for two pairs of probe positions (D89C–T109C and T109C–N140C) with the corresponding experimental data by Lerch *et al*. [83]. First, we calculate the *P*(*r*) of the single states using PDB code 3DMV for the G states and PDB codes 2LCB and 2LC9 for the E states of single and triple mutants, respectively. Second, the *P* (*r*)’s are linearly combined based on the experimentally derived ratios of G and E populations (97:3 for L99A and 4:96 for L99A-G113A-R119P) [82]. Additionally, we predict DEER distance distributions from previously reported metadynamics MD simulations of L99A and L99A-G113A-R119P [80]. In these calculations, the average over the trajectory is weighted by exp (*F*_*bias*_*/k*_*B*_*T*), where *F*_*bias*_ is the final static bias for each frame and *k*_*B*_*T* is the thermal energy. The analysis of a trajectory of 6,670 frames is performed in 11 min on a 1.7 GHz processor executing the following lines of code:

1. `import MDAnalysis`
2. `from DEERPREdict. DEER import DEERpredict`
3. `import numpy as np`
4. `u = MDAnalysis. Universe (’ conf. pdb’,’ traj. xtc’)`
5. `for residues in [[89, 109], [109, 140]]:`
6. `DEER = DEERpredict (u, residues = residues, temperature =298, z_ cutoff = 0. 1)`
7. `DEER. run (weights = np. exp (Fbias /(0. 298 * 8. 3145)))`

In line six, we specify the positions of the spin-labels, the temperature at which the metadynamics simulations were performed and a non-default value for the *Z* cutoff. In line seven, we provide the weights of each trajectory frame, generated from the array of *F*_*bias*_ values.

Figure 3 shows a comparison between the experimental distance distributions obtained by Lerch *et al*. [83] and our predictions. In general, the calculated distributions fall within the experimental ranges of inter-probe distances and are particularly accurate for the D89C–T109C spin-labeled pair in metadynamics simulations. The sharper shape of the experimental *P* (*r*)’s, relative to the calculated distributions, could be due to the cryogenic temperatures at which DEER experiments are conducted, whereas simulations were performed at room temperature. For the T109C–N140C spin-labeled pair of the triple variant, the discrepancy between predicted and calculated *P* (*r*)’s might be explained by considering that distances shorter than 1.5 nm fall below the range probed by DEER experiments. On the other hand, the inaccurate predictions of the T109C–N140C *P* (*r*) for the single (L99A) variant is greater than expected. Such discrepancies may be due both to errors in the protein structure or in the DEER-calculations. While our results cannot distinguish between these scenarios, we follow previous work [14] by examining whether the discrepancies can can be attributed to the error on the Boltzmann probabilities of the rotamer states, 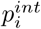. We thus use a Bayesian/maximum entropy procedure to show that a small change in the original rotamer weights can lead to a substantial improvement of the agreement with the experimental data (see Figure S3).

**Fig 3.**
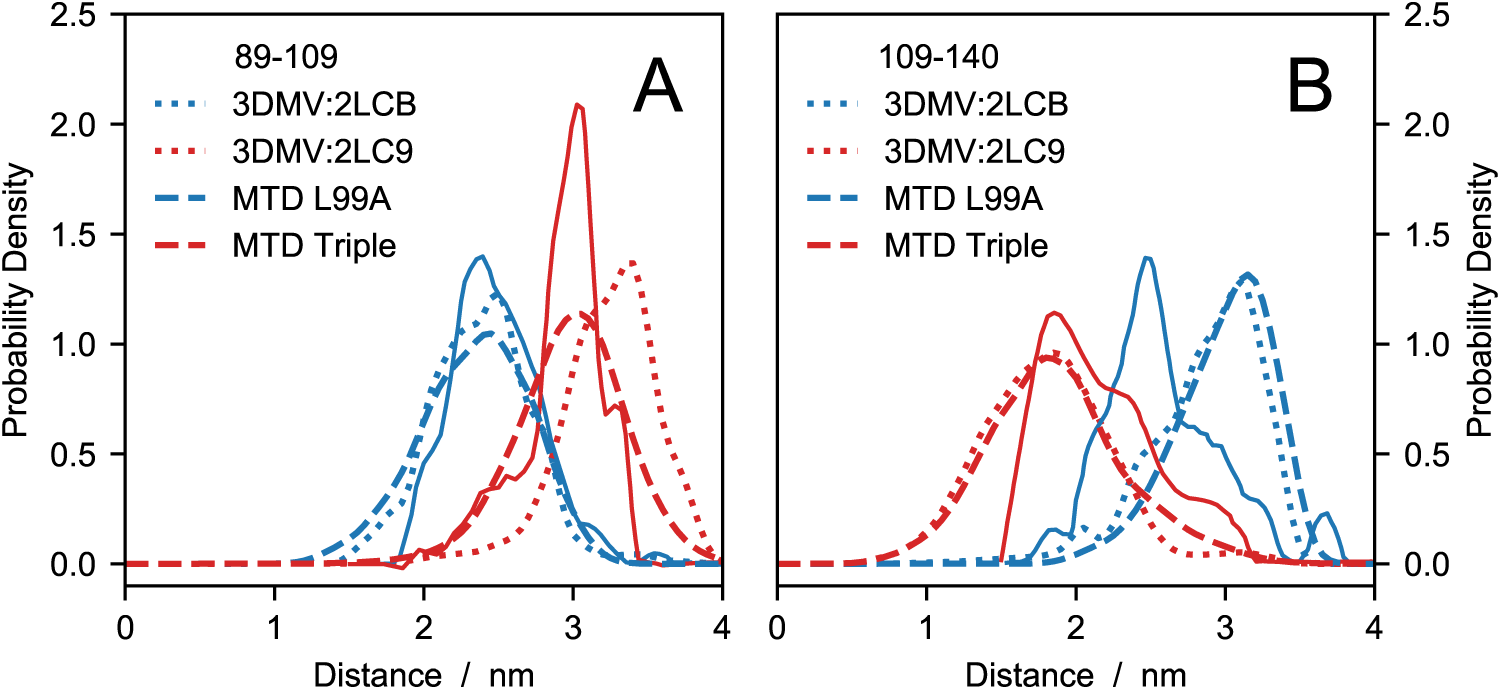
Comparing experiments with simulations and structures of T4 Lysozyme variants. DEER distance distributions for probe positions (A) D89C–T109C and (B) T109C–N140C of the single (blue) and the triple variant (red). Solid lines are the experimental data by Lerch et al. [83], dotted lines are calculated from PDB codes and dashed lines are predictions from metadynamics (MTD) simulations by Wang and coworkers [80].

### Case study 3: PRE data for Acyl-CoA-Binding Protein

The RLA is well known in the EPR community and generally favored over e.g. a C*α*-based approach as discussed elsewhere [3, 13, 26]. In the presented software, we apply the same improved modeling of the probe flexibility also to the prediction of PRE rates and intensity ratios.

Our test data is the PRE data for the bovine Acyl-coenzyme A Binding Protein (ACBP) reported by Teilum *et al*. [53]. In this study the structural behavior of ACBP under native and mildly-denaturing conditions was investigated via the SDSL of five positions in the amino acid the sequence: T17C, V36C, M46C, S65C and I86C. Here, we focus on the native state of ACBP for which an NMR structure comprising 20 conformers has been refined from residual dipolar couplings (RDC) and deposited in the Protein Data Bank (PDB code 1NTI). Figure 4 shows a comparison between the experimental data and the intensity ratios calculated from the Γ_2_ values averaged over the 20 conformations of the PDB entry. A good overall agreement is achieved across the different probe positions. Notably, using the RDC-refined structure, we reproduce most of the structural features observed in the PRE experiments, including the proximity of residues 24, 27, 31 and 34 to the spin-labeled residue 86, which is consistent with a helix-turn-helix motif. The predicted intensity ratios are generated in 2 s on a 1.7 GHz processor executing the following code:

**Fig 4.**
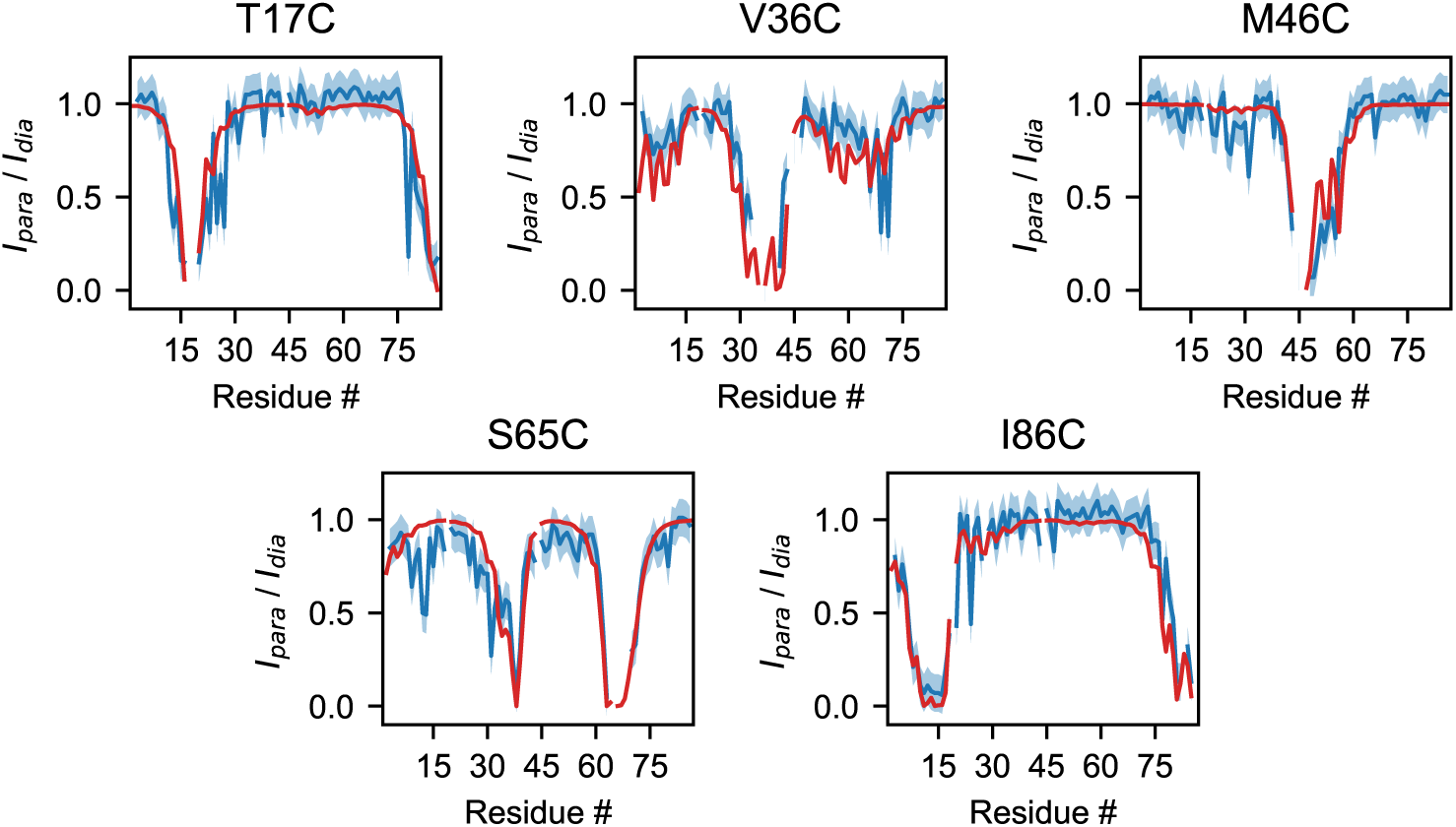
Calculated and experimental PRE HSQC intensity ratios for the T17C, V36C, M46C, S65C and I86C mutants of ACBP. Blue lines represent the experimental data [53], with the associated ±0.1 error shown by the blue shaded areas. Red lines represent intensity ratios calculated from PDB code 1NTI with *τ*_*c*_ = 2 ns, *τ*_*t*_ = 0.2 ns, *t*_*d*_ = 10 ms, *R*_2_ = 12.6 s^−1^.

1. `import MDAnalysis`
2. `from DEERPREdict. PRE import PREpredict`
3. `u = MDAnalysis. Universe (‘1 nti. pdb ‘)`
4. `for res in [17, 36, 46, 65, 86]:`
5. `PRE = PREpredict (u, res, temperature =298, atom_ selection =’ H ‘)`
6. `PRE. run (tau_ c =2 e -09, tau_ t =2*1 e -10, delay =1 e -2, r_2 =12.6, wh = 750)`

At line three, we load PDB code 1NTI as an MDAnalysis Universe object. We then use a for loop to calculate the PRE data from the distances between amide protons and the spin-label N-O group at five different positions along the amino acid sequence. In the last line we specify *τ*_*c*_ = 2 ns, *τ*_*t*_ = 0.2 ns, *t*_*d*_ = 10 ms, *R*_2_ = 12.6 s^−1^ and *ω*_*I*_ = 2*π* × 750 MHz. Per-frame and ensemble-averaged PRE data are automatically saved to files named *res-*\*.*pkl* and *res-*\*.*dat*, respectively, whereas per-frame steric partition functions are saved to *res-Z-*\*.*dat*.

As detailed in Figure S4, the steric partition functions provided by DEER-PREdict can be used to predict whether a position in the sequence is likely to accommodate the paramagnetic probe within the wild-type structure. Besides aiding the interpretation of experimental data, this feature can be instrumental to designing and enhancing the success-rate of time- and labor-intensive SDSL experiments.

As previously discussed, the explicit treatment of the paramagnetic probe may be crucial for the accurate back-calculation of DEER data, and even more so for PRE predictions, due to the ⟨*r*^−6^⟩-dependence of the PRE. A common way to restrain MD simulations or to back-calculate PRE experimental data without explicitly simulating the paramagnetic probe is to approximate the electron location to the position of the C*β* atom of the spin-labeled residue [85]. The advantage of this approach is that (a) multiple labeling sites can be analyzed in a single simulation and (b) the explicit atom is present in the simulation making the calculation of PREs straightforward. C*β*-based calculations may, however, be prone to over- or underestimating electron-proton distances by several Å, thereby introducing a systematic error. The impact of the C*β*-approximation on the accuracy of PRE predictions is illustrated in Figure S5 and Figure S6 for the case of ACBP.

## Conclusion

We have introduced an open-source software solution with a fast implementation of the RLA in tandem with protein ensemble averaging, for the calculation of DEER and PRE data. Using three examples, we have highlighted the capabilities of our implementation: (a) the extension of the RLA for DEER data from a protein ensemble and (b) the calculation of PRE rates and intensity ratios with the same approach.

The structural interpretation of DEER and PRE measurements requires an accurate treatment of the structure and conformational heterogeneity of the spin labels. In the presented software, this is achieved using the RLA and, in the case of the PRE, a model-free approach to describe the dynamics. Relative to simulations of the explicitly spin-labeled mutants, the RLA presents the particular advantage of enabling the prediction for multiple SDSL experiments from a single simulation of the wild type sequence.

## Availability and Future Directions

The software is implemented using the popular trajectory analysis package MDAnalysis, version 1.0 [30] and is available on GitHub at github.com/KULL-Centre/DEERpredict. DEER-PREdict is also distributed as a PyPI package (pypi.org/project/DEERPREdict) and archived on Zenodo (DOI: 10.5281/zenodo.3968394). DEER-PREdict and MDAnalysis are published and distributed under GPL licenses, version 3 and 2, respectively.

DEER-PREdict has a general framework and can be readily extended to encompass non-protein biomolecules as well as additional rotamer libraries of paramagnetic groups. Moreover, the software can be augmented with a module to predict Förster resonance energy transfer data, combining the insertion routines already implemented for MTSSL probes with rotamer libraries for fluorescent dyes.

## Acknowledgments

We thank Robert Best for help with RDC-restrained simulations as well as work on extending DEER-PREdict to use for prediction of FRET experiments. M.B.A.K. acknowledges funding from the Lundbeck Foundation. R.C. acknowledges funding from MINECO (CTQ2016-78636-P). K.L.-L. acknowledges funding via a Sapere Aude Starting Grant from the Danish Council for Independent Research and the Lundbeck Foundation BRAINSTRUC initiative in structural biology.

## Author Contributions

**Conceptualization:** Giulio Tesei, João M. Martins, Micha B. A. Kunze, Kresten Lindorff-Larsen

**Data curation:** Giulio Tesei, João M. Martins, Micha B. A. Kunze, Yong Wang

**Funding acquisition:** Ramon Crehuet, Kresten Lindorff-Larsen

**Software:** Giulio Tesei, João M. Martins, Micha B. A. Kunze, Ramon Crehuet

**Supervision:** Micha B. A. Kunze, Kresten Lindorff-Larsen

**Writing — original draft:** Giulio Tesei, João M. Martins, Micha B. A. Kunze, Kresten Lindorff-Larsen

**Writing — review & editing:** Giulio Tesei, João M. Martins, Micha B. A. Kunze, Yong Wang, Ramon Crehuet, Kresten Lindorff-Larsen

## Supporting Information

### Molecular Dynamics Simulations

#### HIV-1 protease

All simulations were performed using GROMACS 5.1 [86] with the PLUMED 2 [85] plugin. Unbiased and RDC-biased metadynamics [75, 87] simulations where performed starting from a closed conformer after removal of the inhibitor from the X-ray crystal structure (PDB code 3BVB) [88]. The simulated protein is the wild type subtype B from isolate BRU/LAI, which differs from the construct of the reference experiments [44] by the following mutations: M36norleucine, S37N, R41pseudo-homoglutamine, M46norleucine, I63P, I64V, A67*α*-aminobutyric acid and A95*α*-aminobutyric acid. The protein was simulated in a cubic box with a side length of 8.677 nm containing 22,228 water molecules, 59 sodium cations, and 67 chloride anions. Although the K55C mutations have been shown to have a negligible impact on enzymatic activity [64], we maintained the lysine residues at the spin-labeled positions with the aim of capturing the conformational ensemble of the wild type.

The system was equilibrated for 10 ns with a 2-fs time step in the NPT ensemble, using the Berendsen barostat [89] with 0.5-ps coupling constant and isothermal compressibility of 4.5e-5 bar^−1^. Starting from the equilibrated structure, a production run of 500 ns was performed in the NVT ensemble for the unbiased MD simulation. For the restrained simulations, we obtained the backbone N-H RDCs for the inhibitor-free HIV-1PR from Roche *et al*. [66] and applied a linear potential (force constant 25,000 kJ/mol) between the experimental data and the RDCs calculated as averages over 4 independent simulations. Each replica was simulated for 125 ns in the NVT ensemble. We used the AMBER ff99SB*-ILDN force field [90, 91] for all simulations. First, the structure was minimized with the steepest descent algorithm to a tolerance of 10 kJ mol^−1^ nm^−1^ with restrained water molecules. Second, we simulated the system the for 5 ns in the NVT ensemble using the leap-frog integrator with a time step of 1 fs. A 1.2 nm cutoff was used for van der Waals interactions, with a force-switch modification at 1.0 nm. Coulomb interactions were treated with the particle-mesh Ewald method [92] with direct space cutoff of 1.2 nm. The temperature was set to 298 K using the v-rescale thermostat [93] with a 5 ps coupling constant. All the bonds involving hydrogen atoms were constrained using the LINCS algorithm [94].

#### T4 Lysozyme

The simulations of the L99A single mutant and the L99A-G113A-R119P triple mutant of T4 Lysozyme analysed in this study have been reported previously by Wang *et al*. [80]. The X-ray crystal structure of PDB code 3DMV was used as the initial configuration for the G states of both single and triple mutants. Simulations of the E states were started from chemical-shift-derived Rosetta-calculated structures, i.e. PDB code 2LCB and PDB code 2LC9 for single and triple mutant, respectively. Further simulations details can be found in the original article [80].

**Fig. S1.**
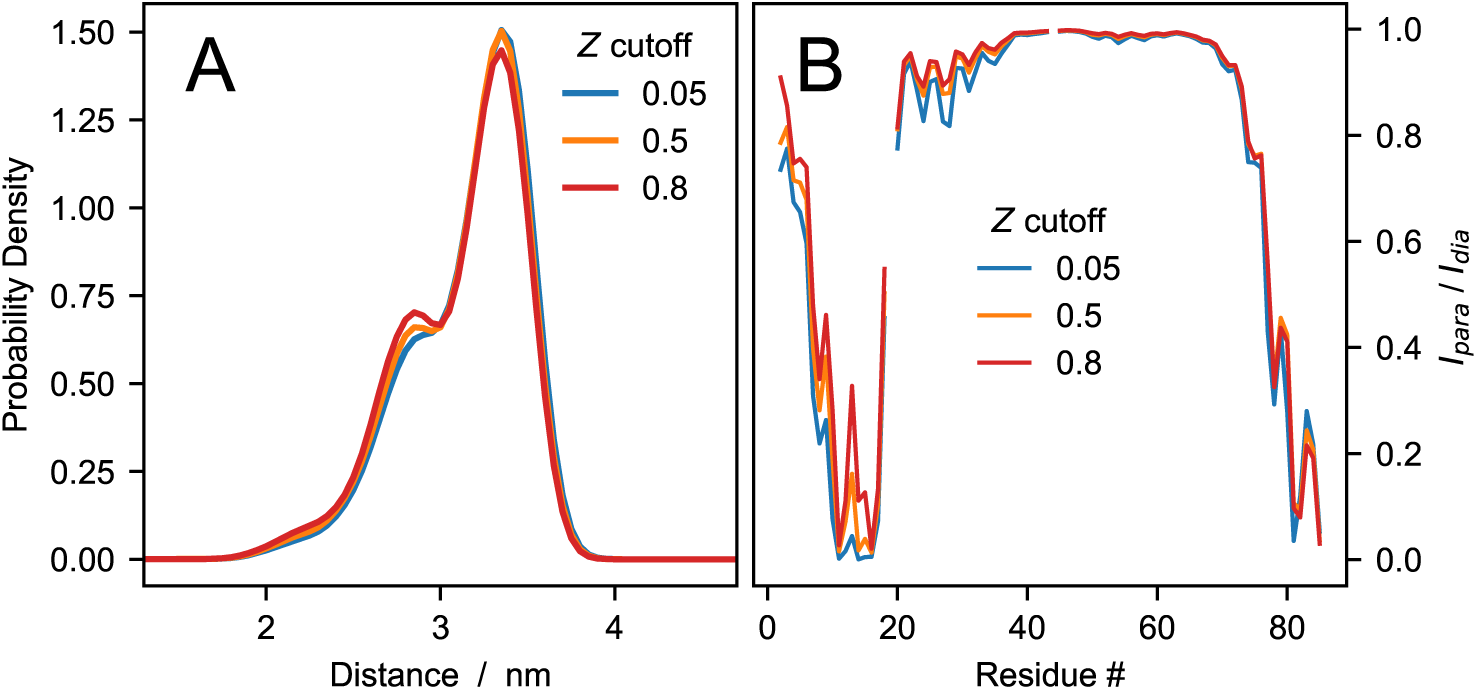
Influence of the *Z* cutoff on predicted DEER and PRE NMR data. (A) DEER distance distributions calculated from RDC ensemble-biased MD simulations of HIV-1PR. (B) Predicted intensity ratios for ACBP spin-labeled at position 86 obtained from PDB code 1NTI with *τ*_*c*_ = 2 ns, *τ*_*t*_ = 0.2 ns, *t*_*d*_ = 10 ms, *R*_2_ = 12.6 s^−1^. DEER and PRE predictions are performed using three different cutoff values of the steric partition function, *Z*, namely 0.05 (blue lines), 0.5 (orange lines) and 0.8 (red lines).

**Fig. S2.**
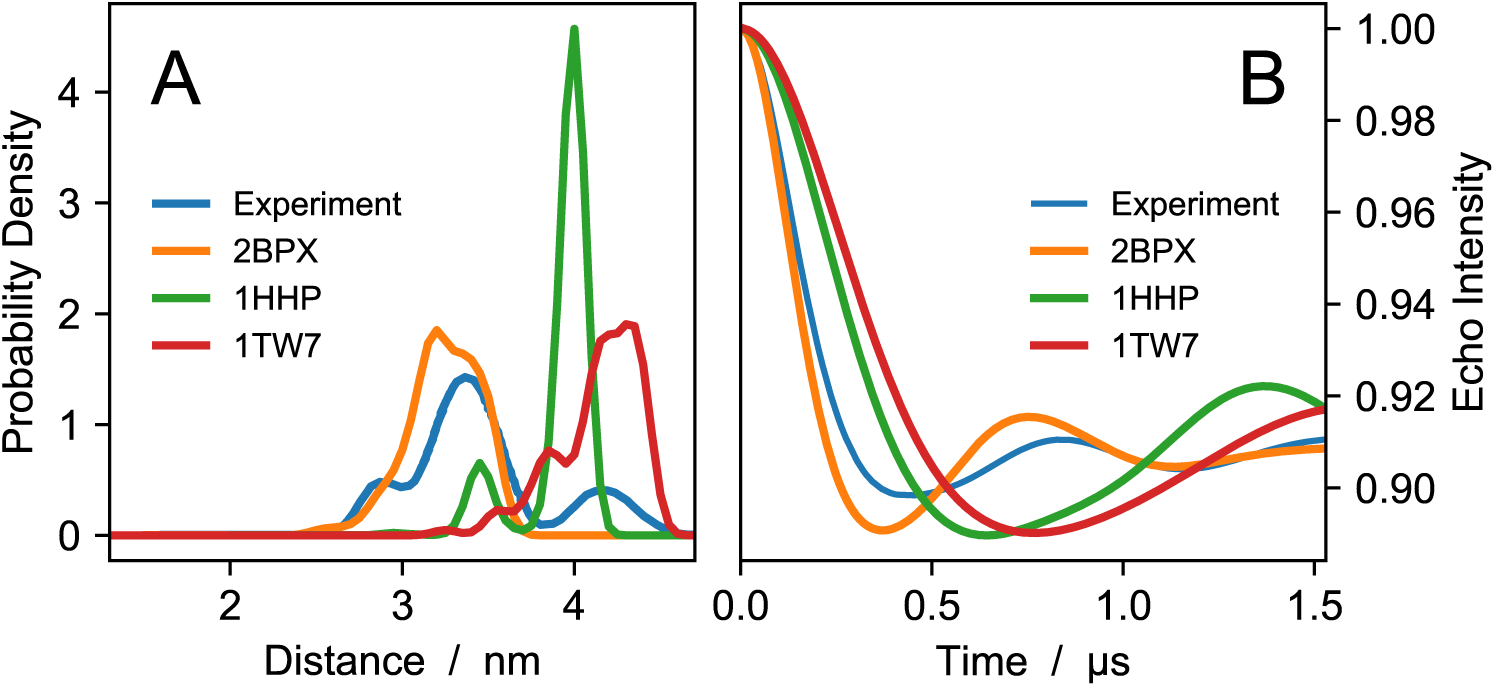
Comparison of DEER data from Torbeev *et al*. [44] with X-ray crystal structures deposited in the Protein Data Bank. This figure shows that although the DEER data is calculated from single X-ray crystal structures, the RLA results in multimodal distance distributions. For example, the K55-K55’ separation between the ammonium groups in 1TW7 is ∼ 3.6 nm, which is consistent with the semi-open conformation, however, the distances between the nitroxide groups of the MTSSL conformers span a wide range between 3.3 and 4.4 nm. DEER distance distributions (A) and echo intensity curves (B) obtained by Torbeev *et al*. [44] from DEER experiments (blue), and calculated using X-ray crystal structures representative of closed (PDB code 2BPX, orange), semi-open (PDB code 1HHP, green) and wide-open (PDB code 1TW7, red) HIV-1PR conformations.

**Fig. S3.**
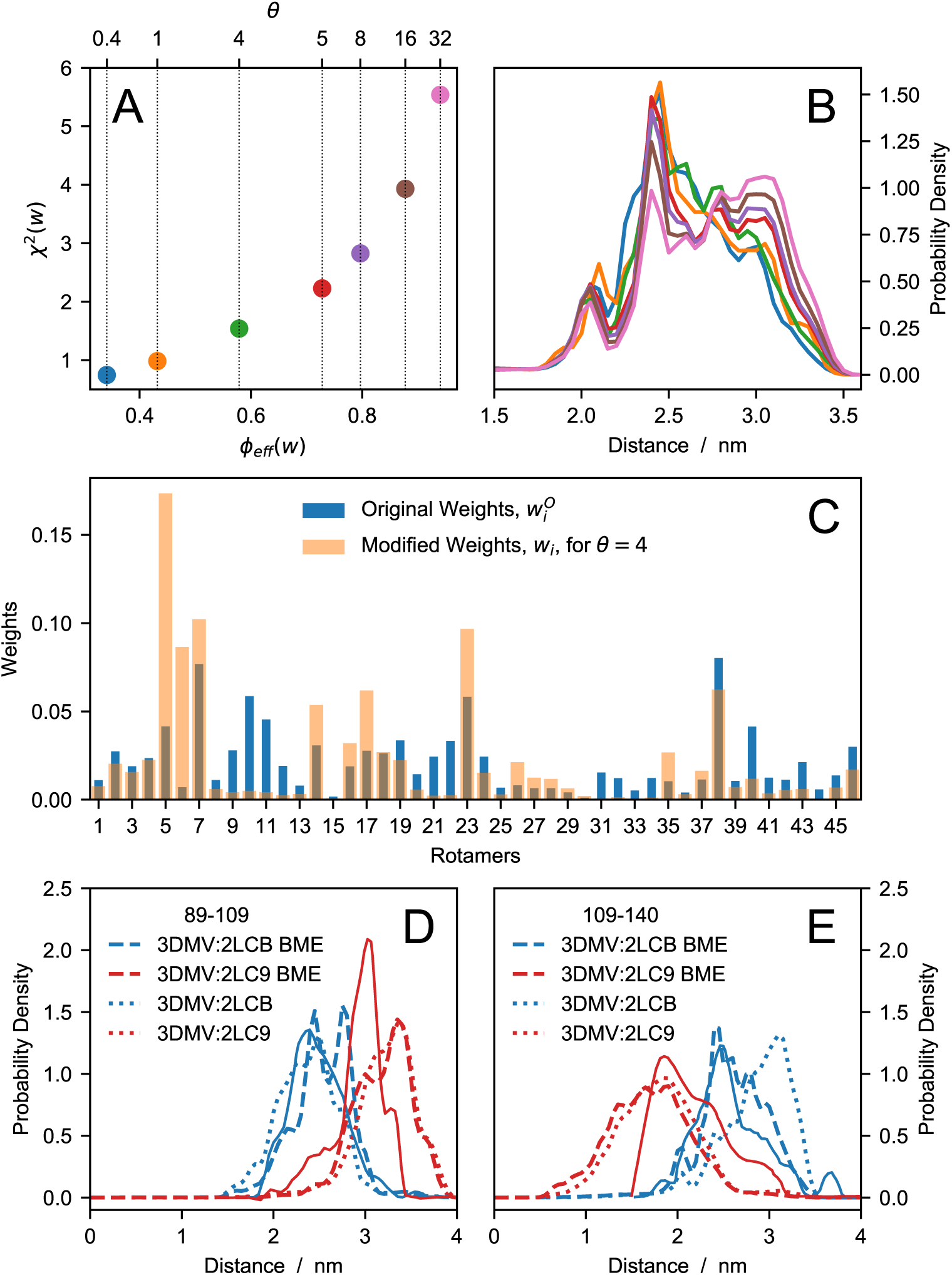
Optimization of rotamer weights using a Bayesian/maximum entropy procedure. We use a Bayesian/maximum entropy (BME) procedure to find a modified set of rotamer weights for the MTSSL 175 K rotamer library [24], ***w***, which improves the agreement between predicted and experimental T109C–N140C *P* (*r*)’s for the single variant of T4L. The prediction is based on the 97:3 linear combination of *P* (*r*)’s from PDB codes 3DMV and 2LCB (corresponding to the populations of these two states) whereas the experimental data is from Lerch *et al*. [83]. Simulated annealing is used to minimize the cost function *ℒ* (***w***) = *χ*^2^(***w***) − *θS*(***w***) where *χ*^2^(***w***) is the sum of the squared differences between predicted and experimental *P* (*r*)’s, *θ* quantifies the confidence in the original weights, ***w***^***O***^, and 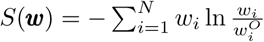 is the relative entropy defined as the negative of the Kullback-Leibler divergence between the modified weights of the *N* = 46 rotamers, *w*_*i*_, and the initial 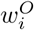. The effective fraction of rotamers used in the reweighted ensemble, compared to the original library, is quantified as *ϕ*_*eff*_ (*w*) = exp [*S*(*w*)]. We scan various values of *θ* and select the weights obtained using *θ* = 4 as the smallest modification resulting in a substantial decrease in *χ*^2^. It is noteworthy that the change in weights has a lesser impact on the triple variant than on the single variant. Moreover, albeit being optimized against the T109C–N140C *P* (*r*) of the single variant, the modified weights lead to an overall improvement in accuracy, with the average *χ*^2^ over the four *P* (*r*)’s decreasing from 7.5 to 5.9. (A) *χ*^2^ vs *φ*_*eff*_ for various values of the confidence parameter, *θ*. (B) Distance distributions calculated from PDB codes 3DMV and 2LCB, using optimized weights obtained for various *θ* values. (C) Original [24] and modified weights of the MTSSL 175 K rotamer library after BME reweighting with *θ* = 4. DEER distance distributions for probe positions (D) D89C–T109C and (E) T109C–N140C of the single (blue) and the triple variant (red). Solid lines are the experimental data by Lerch *et al*. [83]; dotted and dashed lines are from PDB codes 3DMV, 2LC9 and 2LCB using the original and the BME-reweighted (*θ* = 4) MTSSL 175 K rotamer library.

**Fig. S4.**
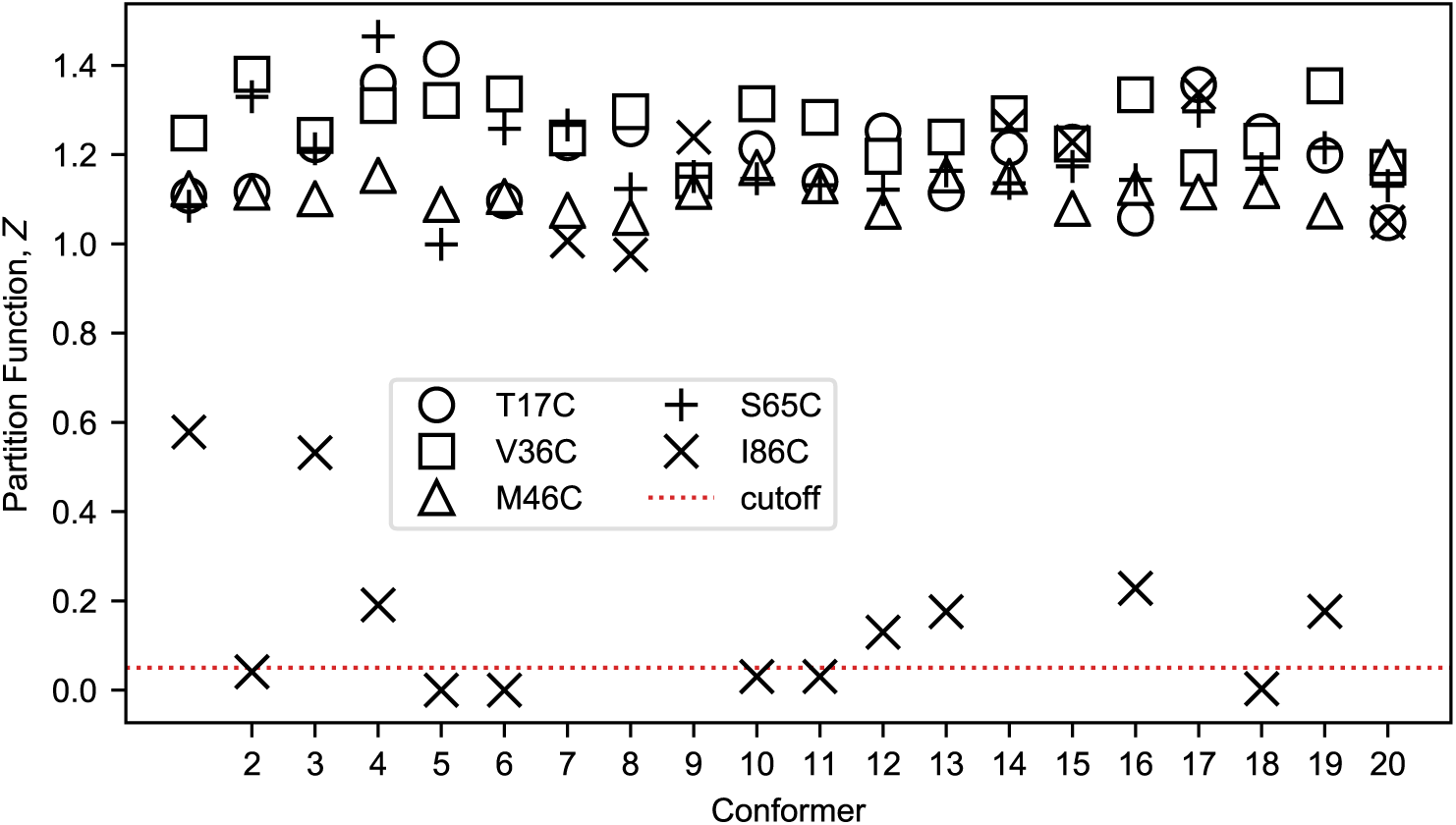
Steric partition function quantifying the fitness of the rotamers at the spin-labeled site. While for most spin-label sites of ACBP the steric partition function, *Z*, varies between 1 and 1.5, for the placement of the probe at residue 86, *Z* drops below the cutoff of 0.05 proposed by Polyhach *et al*. [24] in five conformers out of 20. The significantly lower *Z* values indicate that residue 86 is in a tightly packed region in the protein structure and that spin-labeling position 86 can lead to structural deformations or changes in the populations of the conformational ensemble. This observation is consistent with stability experiments performed by Teilum *et al*. on wild type and spin-labeled mutants [53], showing that I86C is the least stable of the studied spin-labeled mutants. Although the RLA assumes that the overall protein conformation is unaffected by the presence of the spin-label, the case of the I86C mutant of ACBP highlights how DEER-PREdict can help to identify spin-labeled sites that are likely to violate this assumption in folded proteins. Steric partition function calculated from rotamer-protein van der Waals interactions for five spin-labeled mutants of ACBP. The horizontal dashed line indicates the cutoff used in the criterion for discarding protein conformations where the placement of the rotamer is characterized by steric clashes with the surrounding residues.

**Fig. S5.**
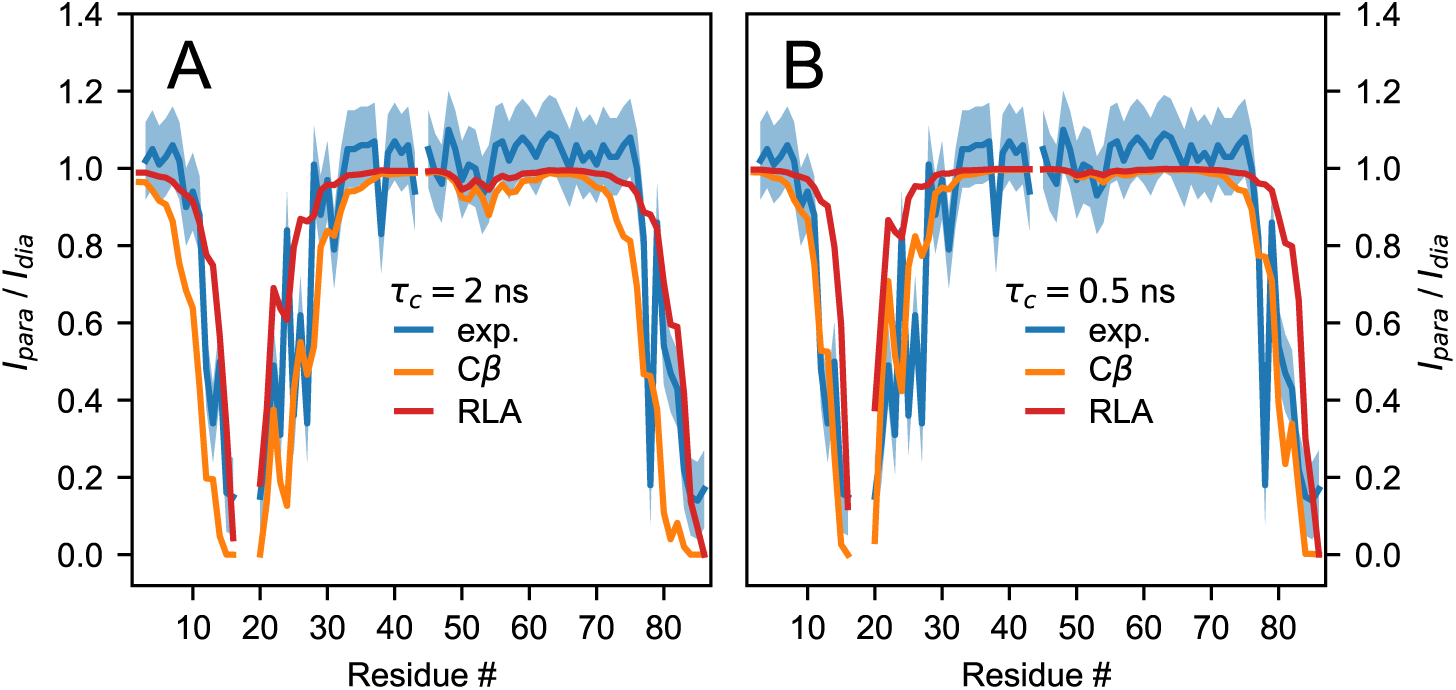
Comparison with C*β*-based PRE Predictions. The C*β* approximation may overestimate the effect of transient tertiary contacts on the PRE rates of ACBP, yielding experimentally consistent predictions only when the time scales for the reorientational dynamics is artificially made fast. PRE intensity ratios for ACBP spin labeled at position 65 calculated for (A) *τ*_*c*_ = 2 ns and (B) *τ*_*c*_ = 0.5 ns. Blue lines represent the experimental data [53], with the associated ±0.1 error shown by the blue shaded areas. Orange and red lines represent C*β*-based and RLA-based predictions, respectively.

**Fig. S6.**
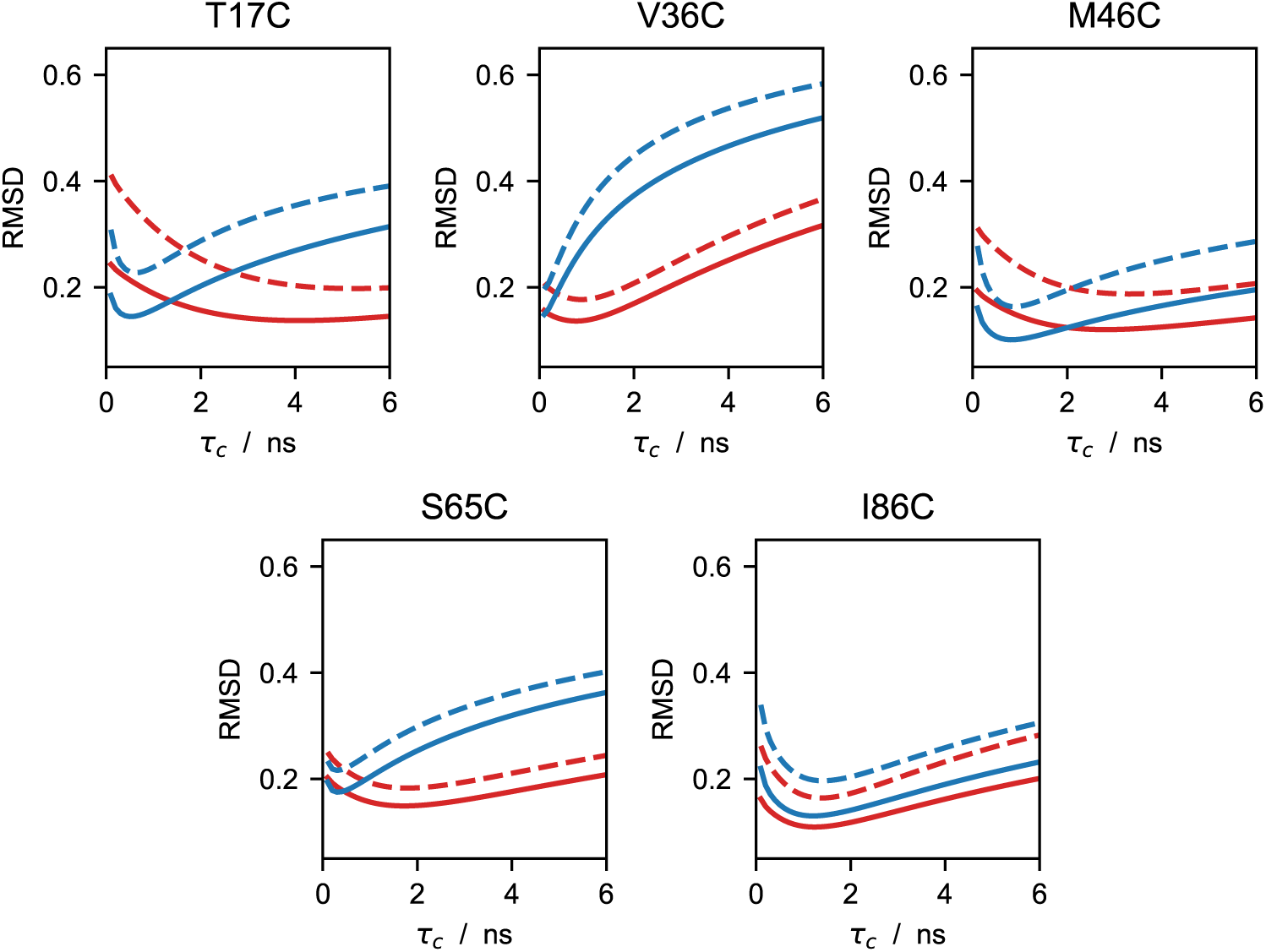
Comparison of optimal *τ*_*c*_ values for RLA vs. C*β*-based approach. This figure illustrates the systematic time-scale difference when using explicit MTSSL probes instead of approximating the location of the unpaired electron with the position of the C*β* atom of the spin-labeled residue. Compared with the *C*_*β*_-based approach, the RLA improves the accuracy in reproducing the experimental PRE data, as shown by the generally lower RMSD values. When using the RLA, the *τ*_*c*_ values that minimise the RMSD are found in the range of typical reorientational correlation time constants for proteins of ∼ 100 residues (2–5 ns), and closer to the experimentally-derived value of 4 ns [53], whereas the C*β*-based approach underestimates the optimal *τ*_*c*_. Dependence on *τ*_*c*_ of the RMSD between experimental and predicted PRE ratios of ACBP. Red and blue lines are obtained using the RLA and approximating the electron location with the position of the C*β* atom, respectively. Solid and dashed lines represent the RMSD values calculated from all the data points and from intensity ratios in the dynamic range 0.1 < *I*_*para*_ / *I*_*dia*_ < 0.9.

## Notes

https://github.com/KULL-Centre/DEERpredict

### Competing Interest Statement

The authors have declared no competing interest.

### Summary of Updates

Minor revisions of text and new analysis with reweighting of rotamers (Fig. S3)

